# Assessing wetland conservation action effectiveness in the Mekong Delta with Sentinel-1 Synthetic Aperture Radar data

**DOI:** 10.1101/2025.07.14.664714

**Authors:** Sietse O. Los, Saran Prak, Bena R. Smith, Olly van Biervliet, Geoff M. Hilton, Daphne Kerhoas

**Affiliations:** Conservation Evidence, WWT, Slimbridge, Gloucestershire, GL2 7BT, UK; WWT Cambodia, 17B, Street 494, Sangkat Phsa Deoum Thkov, Khan Chamkarmon, 12310 Phnom Penh, Cambodia; International Programmes, WWT, Slimbridge, Gloucestershire, GL2 7BT, UK

## Abstract

Concerns about the effectiveness of nature conservation efforts have led to the adoption of evidence-based approaches and more rigorous evaluation methods. This is especially relevant to wetlands, which are crucial for ecosystem services and face global threats from human actions. In the present study, 10-m Sentinel-1 synthetic aperture radar (SAR) data, which provide global coverage regardless of cloud cover, are used to assess the effectiveness of wetland restoration activities which aim to promote wetter conditions. These were carried out during 2022-2024 in the Boeung Prek Lapouv area (Mekong delta, Cambodia). A method was developed to separate SAR signals from the dry and wet seasons to infer variations in groundwater level and wetness retention. Analysis of these signals showed that interventions were effective to varying degrees with the construction of new pools, the lowering of grounds and blocking of ditches being the most impactful.

## 1. Introduction

Concern over ineffective nature-restoration practices led to the promotion of adaptive management and evidence-based conservation approaches (Bennett 2016). Interventions are increasingly evaluated with scientifically rigorous designs to improve conservation management (Hoffman et al. 2010, Pressey et al. 2021). Evaluation methods include literature reviews and meta-analyses (Oldekop et al. 2015), analysis of trends in IUCN red-listed species (Brooks et al. 2009, Fernandez et al. 2022) and of stakeholders’ perceptions (Ferraro and Pressey 2015, Bennett 2016).

In the present study, Sentinel-1 synthetic aperture radar (SAR) data are used to assess the effectiveness of wetland restoration techniques in the Boeung Prek Lapouv area (Mekong Delta, Cambodia). SAR data are increasingly used for wetland monitoring since field monitoring is expensive, time consuming, sometimes unsafe and areas can be remote and difficult to access. Most SAR studies are applied to wetlands in the Americas and China (Adeli et al. 2020). In Cambodia, SAR data are used to monitor the extent of waterbodies including flooded areas (Sabel et al. 2015) and to estimate land-cover type (e.g. rice paddies, Fernández-Urrutia et al. 2023).

The Mekong Delta is known as the rice bowl of South-east Asia due to the annual output of 25 million tons of rice (Eyler 2019), while also being very important for wetland biodiversity (Campbell 2012). Despite this importance to people and wildlife, it’s facing numerous threats (Eyler 2019). Of particular concern are increasingly drier conditions from January until July (dry season) at Boeung Prek Lapouv which are caused by the construction of canals (Yang et al. 2025) and the extraction of groundwater for domestic and industrial use (Minderhoud et al. 2017). In addition, the seasonality of flooding and recession of the Mekong is expected to be sensitive to climate change (Khanal et al. 2021). Water management and the rapid development of agriculture over the past 50 years have led to the disappearance of much of the natural wetland habitat in the Cambodia part of the Mekong Delta (WWT 2023, Indo-Burma Ramsar Regional Initiative 2022). Remaining natural wetlands support several threatened species, including the iconic and globally threatened Sarus crane (*Antigone antigone*, Birdlife International 2016), the tallest flying bird in the world, which spends the non-breeding season (December – February) in the area.

Our study has two aims: the first is to assess the effectiveness of conservation activities in the Boeung Prek Lapouv area to restore wet conditions with the purpose of reverting current dry vegetation communities to wetter ones and promote thriving wildlife, in particular the Sarus crane. Identifying the most effective interventions will also inform future management of wetlands in the area. The second aim is to investigate the usefulness of Sentinel-1 10-m synthetic aperture radar (SAR) data (Section 3.3) for the assessment of the effectiveness of management interventions. This use of SAR data, studying the impact of conservation management in the Mekong Delta, is novel. A challenging aspect of this study is the separation of SAR signals from the wet and dry season, which show similar behaviour (Bangira et al. 2021). Separation of these signals allows both monitoring of flooding and of variations in soil moisture.

## 2. Study area

The Boeung Prek Lapouv Protected Landscape (0.3 to 4.8 m above MSL) is situated in the Mekong Delta (Cambodia, midpoint 10.75° N 105.02° E; Fig. 1) in the western floodplain of the Bassac River and encompasses 8,305 ha of wetland. It obtained ‘Management and Conservation Area for Sarus crane and other birds’ status in 2007 and ‘Protected Landscape’ status in 2016 (equivalent to an IUCN category IV protected area). Land cover consists of rice agriculture (62.7%), grassland (29.6%), forest and scrub (6.2%), and open water (1.2%). About 22 villages (∼12,000 people) use the wetland to farm rice and to collect natural resources including fish, edible plants, firewood and grass (Sophanna et al. 2019). Water levels rise in Boeung Prek Lapouv at the onset of the rainy season from their lowest levels in late May and flood the entire site around October (3-4 m depth, Fig. 1.b) followed by a rapid recede until December and slow recede until May. Soils, in some areas containing acid sulphate horizons, are formed of young alluvial clay and silty-clay swampy-marine sediments interchanged with alluvial tidal-flat deposits (Seng Kim Hout 2004).

**Figure 1:**
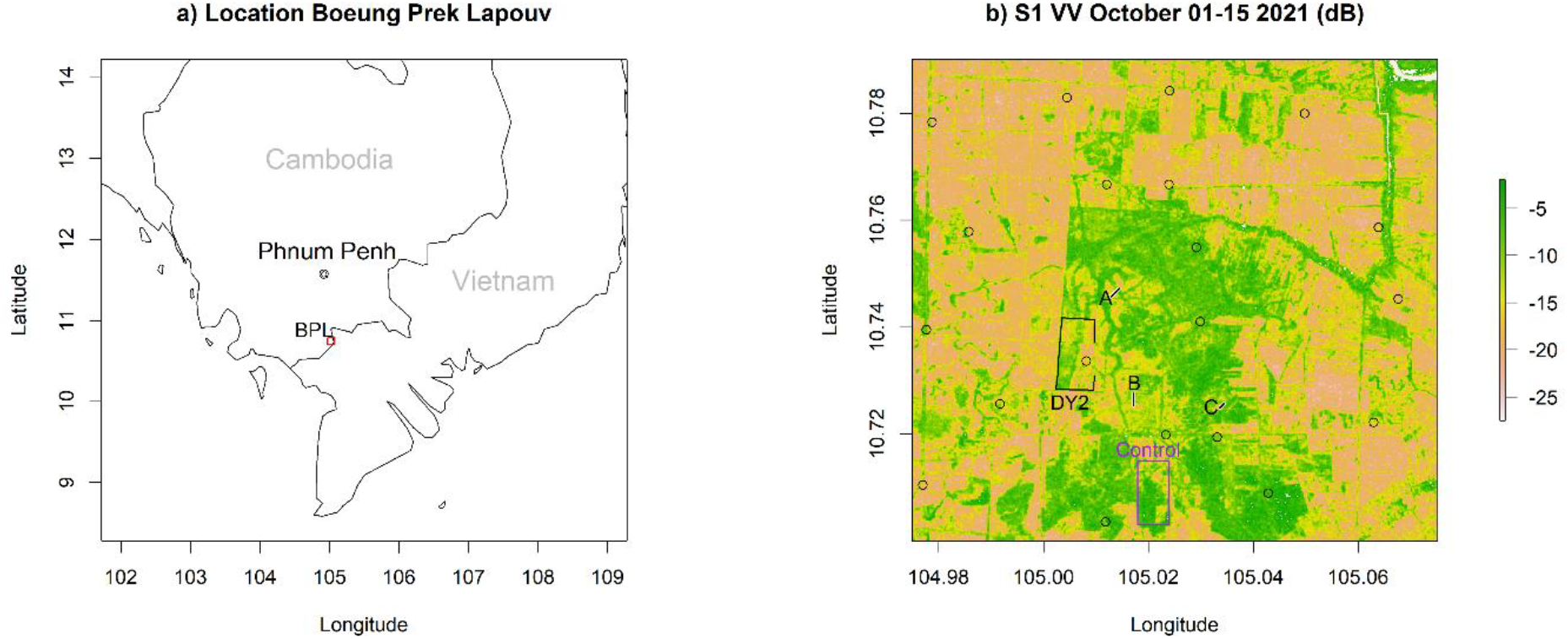
a) Location of the study area marked by the rectangle (BPL=Boeung Prek Lapouv). b) Sentinel-1 VV polarization image of the wet season showing locations of dipwells (circles) and of selected interventions (Section 5): A-B mark transects with filled ditches and C a transect with an unfilled ditch (control), DY2 was enclosed in 2023 and “Control” (no activities) was used for comparison.

In recent years, wet season floods in Boeung Prek Lapouv receded earlier and desiccation at the site during the dry season became more severe, with negative impacts on Sarus crane and other wetland species. Consequently, conservation interventions to retain water for longer into the wet season were trialled. Activities between 2022-2024 comprised: (1) blocking of ditches (by filling with soil) to reduce draining (N=7, total length=1.8 km); (2) the construction of ephemeral pools (N=10, total area=18.9 ha); (3) the enclosure of a large rectangular area with a dyke to retain flood water (N=1, area=108 ha) and (4) ground lowered (N=7, total area=53.4 ha) by removal of 10 cm of soil thus exposing wetter soil situated closer to the groundwater table. In 2024 excavated soil was used to build mini bunds around the pools and lowered grounds to improve retention of water. Sites were selected based on topography, evidence of drying out (shrub encroachment) in consultation with stakeholders and government authorities. Areas with disputed land ownership, sensitive wildlife habitat or iron-sulphide soil layers were avoided.

## 3. Data

### 3.1 Groundwater levels

Groundwater levels were measured at weekly intervals during the dry seasons (January until July) of 2021–2024 in 20 dipwells of about 2 m depth (Fig. 2). Dipwells were flooded during the wet season (August–December)

**Figure 2:**
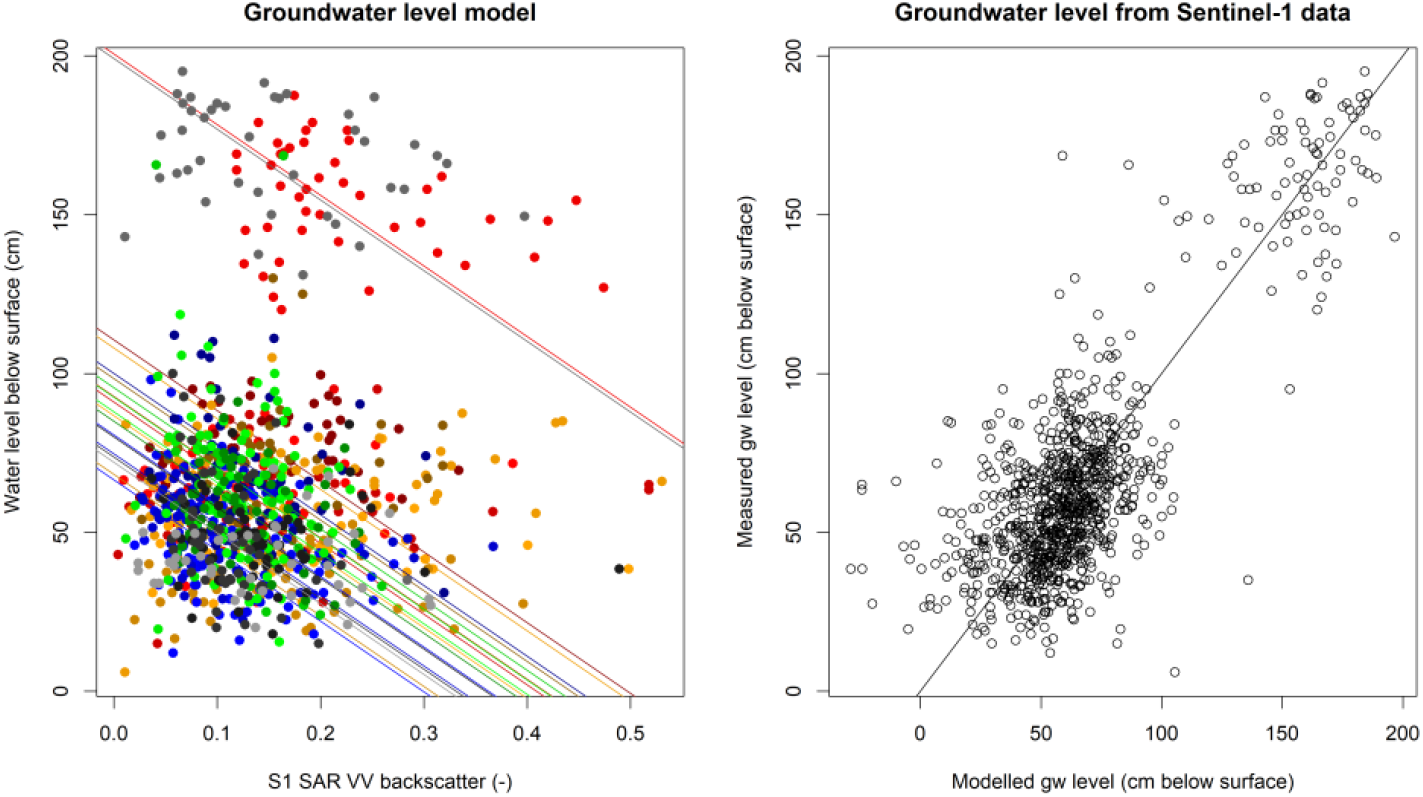
Comparison of Sentinel-1 VV backscatter (dimensionless) with groundwater levels (cm below the surface; 2021-2024). The coloured lines (left panel) show fits for each of the 20 dipwells (Fig. 1). The right panel shows modelled versus observed groundwater level (r = 0.81; RMSE = 21 cm).

### 3.2 DEM

A UAV was flown in July 2021 to obtain a digital elevation model (DEM) represented by 10 cm contours at 2 cm (centre of BPL) and 5 cm (perimeter) resolution (Sivuth 2021). Data were sampled to 30 m resolution.

### 3.3 Sentinel-1 data

The Sentinel-1A (January 2021 – May 2025) 1B (2021) and 1C (2025) Ground Range Detected (GRD) Synthetic Aperture Radar (SAR) data were obtained from the Copernicus Data Space Ecosystem (Copernicus Sentinel data 2025). This product contained vertical-vertical polarized (VV) and vertical-horizontal polarized (VH) C-band data (5.405GHz) at approximately 10 m resolution. Corrections were applied with the SeNtinel Applications Platform (SNAP) toolbox for satellite orbit variations, thermal noise, calibration, terrain flattening and terrain geometry (SNAP 2024). Data were cropped to the research area (Fig. 1) and individual images were averaged to obtain twice-monthly composites (days 1–15 and remaining days of the month).

### 3.4 Rainfall re-analysis

ERA5-Land 9 km monthly rainfall reanalysis (Muñoz-Sabater et al. 2021) for January 2021 until September 2024 was obtained from the Copernicus Climate Change Service (C3S, 2024). The reanalysis was used to indicate the start of the dry and wet seasons (Section 4.2).

## 4. Methods

### 4.1 Groundwater-level variations from SAR data

Variations in groundwater levels (distance below the surface) for 2021–2024 were estimated from SAR data using a regression analysis, which is a common approach for local analysis (Kornelsen and Coulibaly, 2013). The groundwater levels were averaged over the twice-monthly periods and compared with the SAR-VV data (Fig. 2). A regression line was estimated with canonical correlation analysis (CCA; González and Déjean 2023, R Core Team 2024). CCA was preferred over ordinary least squares because it is resistant to errors in the independent variable (SAR data).

It was assumed that the slope of the relationship between groundwater levels and SAR-VV data was the same for all dipwells and that the offset was location dependent:

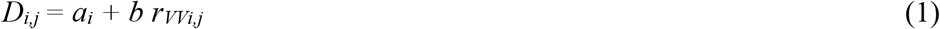

With *D* being the depth of the groundwater table, subscript *i* indicating one of twenty dipwells, subscript *j* the time of observation, *a*_*i*_ the offset for dipwell *i, b=*-222.3 the slope of the regression line, and *r*_*VV*_ the VV backscatter signal. The coefficient *b* allows conversion of the VV signal into a relative change in groundwater level (Section 5). Only seasonal variations in groundwater level were estimated; absolute groundwater levels were not estimated. Note that the largest negative values are closest to the surface.

### 4.2 Flood estimation with SAR data

The analysis of BPL with SAR data is complicated by alternating dry and flooded conditions; the Sentinel-1 VV and VH signals are highest for near-saturated soils but decrease both for drier and wetter (flooded) conditions (Bangira et al. 2024). This response is illustrated in Fig. 3: for dry conditions both VV and VH are low (A) and both increase in response to higher soil moisture (B). When the soil becomes saturated and a thin layer of water develops at the surface, the VH signal decreases but the VV increases because it is reflected off the water surface on to the vegetation and back into the sensor (C – double bounce). When flooding continues and vegetation drowns, both VV and VH become low; usually lower than for a dry, bare soil (D). SAR data from flooded and dry conditions were separated by first identifying cycles consisting of a decrease and subsequent increase in VV, VH or the ratio between them, VH / VV (all converted to dB). The cycle is attributed to a flooding event if very low values occur during the cycle that are associated with open water (VV < -15.5 dB; VH < -24.5 dB; Fig. S1), or if rainfall is high at its start; otherwise, the cycle is attributed to a dry event. The Supplement provides details of the algorithm.

**Figure 3:**
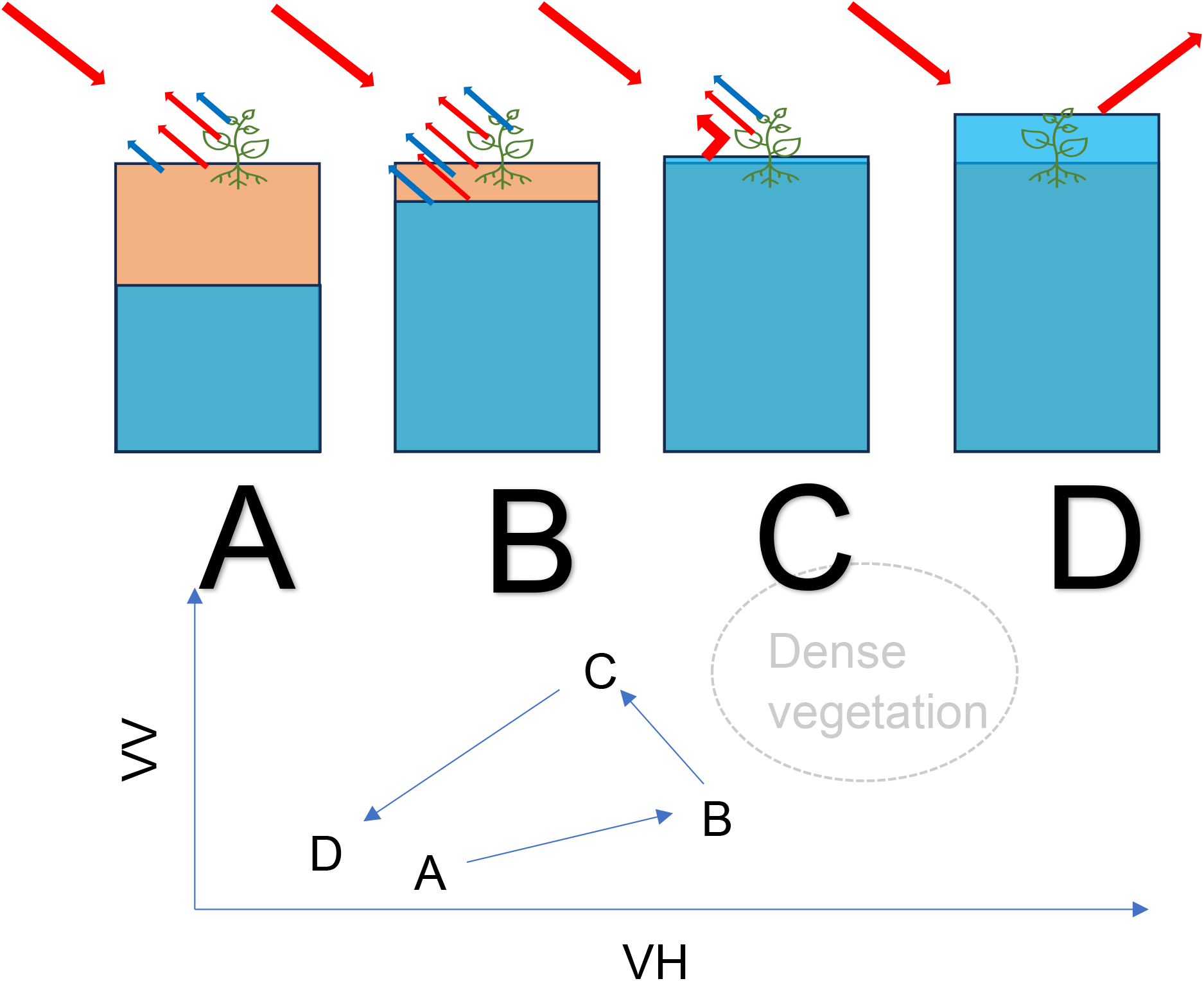
SAR C band response (VV and VH) for different soil moisture conditions and open, short vegetation. A: dry soil and vegetation. B: wet soil and vegetation. C: Wet surface below vegetation. D: Flooded vegetation.

## 5. Results

The impacts of the four types of intervention conducted between 2022 and 2024 (Section 2) were assessed from the estimated groundwater-level variations and flooding occurrences (Section 4.2). The analysis discussed with detailed examples (Section 5.1 – blocked ditches; Section 5.2 – enclosed area) and is summarised in section 5.3 for all interventions.

### 5.1 Blocked channels

Fig. 4 illustrates the analysis of ditches blocked in 2024 (200 m transects A and B, Fig. 1) and of an unblocked ditch (100 m transect C). SAR VV and VH signals were averaged over a rectangular area that extended 10 m either side of a ditch. During the dry season the VV signal is linked to variations in groundwater level, and the VH signal to moisture variations in the vegetation canopy. For the blocked channels A and B, the decline in VV during the 2024 dry season compares to that in previous years (Fig. 4) indicating similar groundwater levels. This contrasted with conditions for the unblocked ditch at transect C where the VV signal was lower indicating lower groundwater levels and drier conditions in 2024. Since the drying out was not observed at the blocked ditches, the intervention was deemed effective.

**Figure 4:**
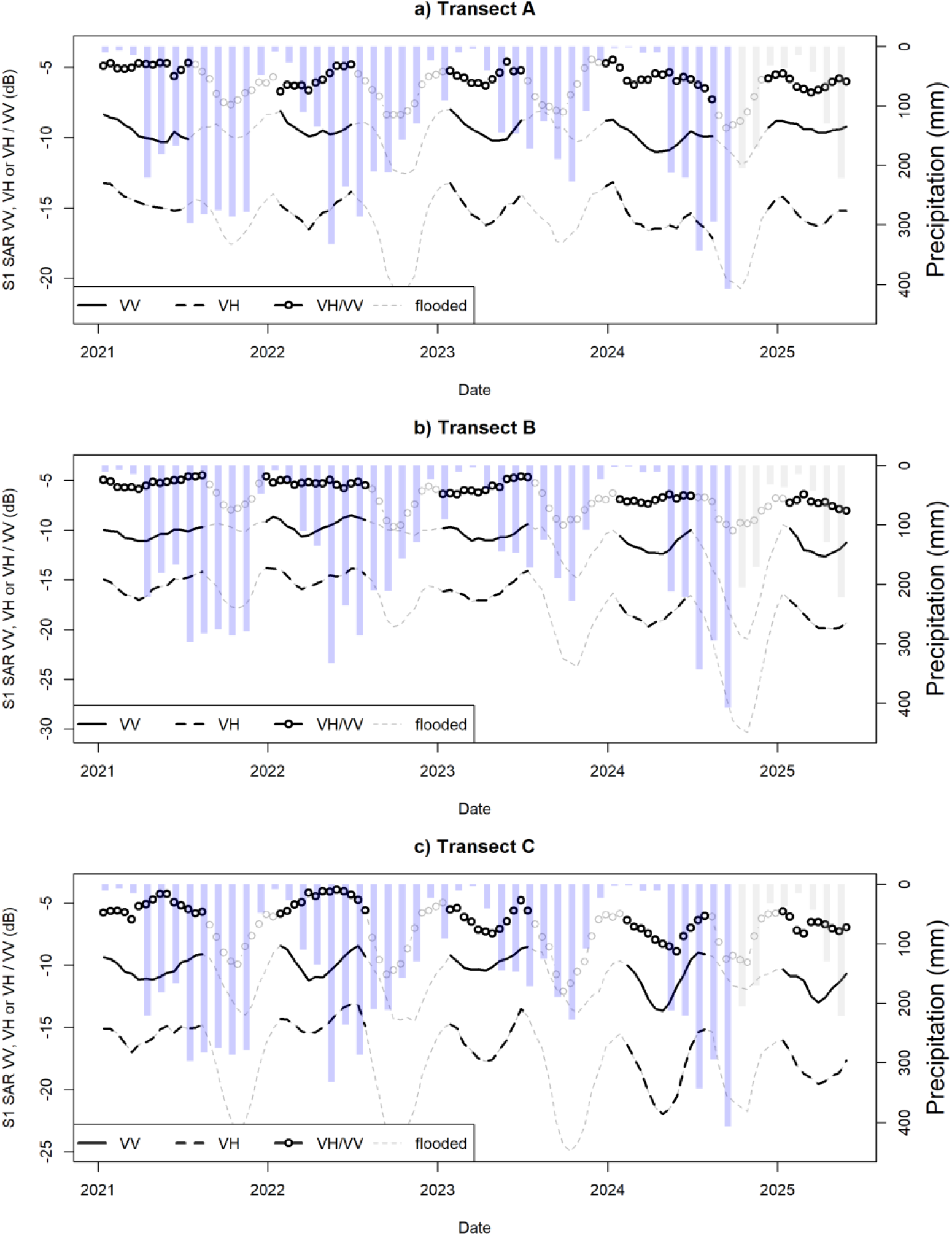
a) Smoothed (2.5 month moving average) Sentinel-1 SAR VV, VH and VH/VV (dB) time series for transect A (blocked ditch; location in Fig. 1.b). Black lines show groundwater variations during the dry season (B – A – B trajectory in Fig. 3) and grey lines flooding during the wet season (B – C – D – C – B in Fig. 3). The bars indicate precipitation (purple bars: monthly values; grey bars: multi-year average monthly values). The effect of a double bounce, marked by a decrease in VH and increase in VV (Section 4.2), can be seen at the start of the wet season in 2021 – 2023 (a and b). b) and c) same as a) but for Transect B and C, respectively. Transect C (no ditch blocking) is drier during the 2024 and 2025 dry seasons than the other two transects.

### 5.2 Enclosed area

Changes in dry-season groundwater levels for the enclosed area (Fig. 2) was estimated by converting SAR VV data to groundwater-level variations using the positive value of the slope from eq. 1. For each pixel, near-saturated conditions (B in Fig. 3) were assumed to correspond to the 1 % wettest value (in cm) of the dry season. Then, for saturated conditions (B-C-D in Fig. 3) the wetness was estimated by adding 10 cm to the 1% value – this is approximately the minimum distance to the surface measured in the dipwells (Fig. 2). These adjusted levels are expected to be near the surface. Another 10 cm was added if the VV was below -15.5 dB or the VH below -24.5 dB; it was assumed that at this level most of the short vegetation was drowned. Unlike the VV data, the new time series (Fig. 5) have a uni-directional relationship with wetness. The estimates of flooded area (bottom panel Fig. 5) show that in 2024, the year after the intervention, both the start and end of flooding are delayed compared to 2022. Groundwater levels (top panel Fig. 5; adjusted to show the same direction as the SAR-derived water-level variations) showed that the enclosed area was wetter in the 2024 dry season compared to 2022; by contrast, conditions during the dry season were of 2025 did not show an increase in wetness compared to 2022.

**Figure 5:**
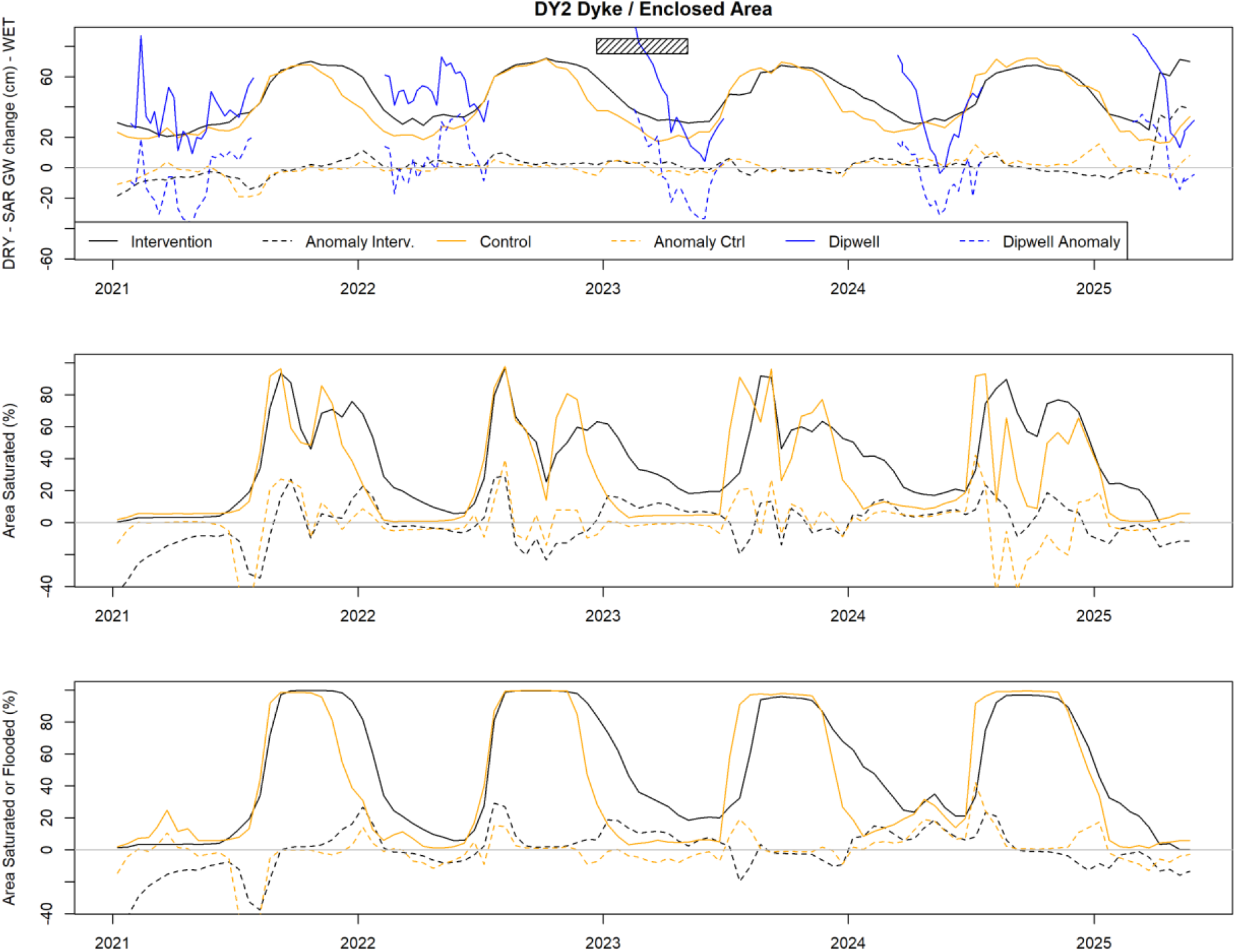
SAR estimated groundwater level variations (top), saturated area (middle) and sum of flooded and saturated area for the DY2 enclosed area constructed in 2023 (Location in Fig. 2; timing indicated by hatched bar). Also shown are anomaly time series (departures from 2-weekly mean values – dashed lines) to aid assessment of wetter or drier conditions for a particular time of year. Dipwell data show the same direction of change as the SAR estimated groundwater-level variations. Effect of pools constructed within the enclosed area are not included. The effect of the intervention for Feb – April 2024 and 2025 appears small in the SAR data.

### 5.3 Combined results of interventions

The analysis (Section 5.2) was applied to all interventions (pools, lowering of the ground, enclosed area and ditch filling; see Section 2.) carried out in 2022–2024. The differences in February to April SAR-derived groundwater variations for the years before and after the interventions were calculated and the same differences for the control were subtracted to obtain an indication of change in wetness (Fig. 6). For ditches, groundwater variations from a rectangle 10 m either side of the ditch were averaged and were compared against the control ditch (transect C); for other interventions, only the areas affected were considered and were compared to changes in the rectangular control site (Fig. 1). The construction of pools in 2024 resulted in the largest increase in wetness of all the interventions (Fig. 6). The next largest effect was found for lowering of the ground (2024) and for ditch filling; the enclosure of an area by a dyke led to wetter conditions in 2024 but not in 2025 (Fig. 5).

**Figure 6:**
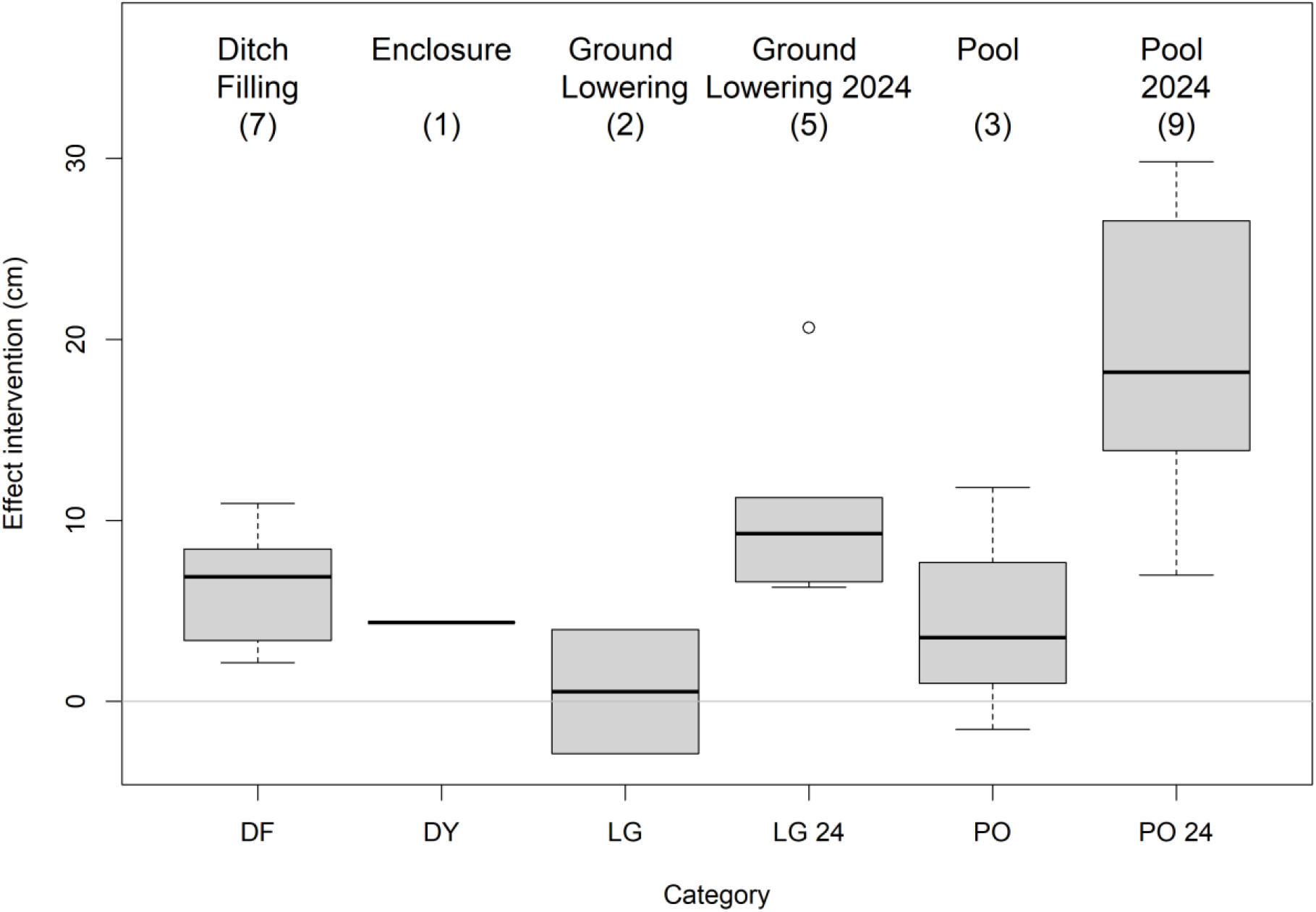
Boxplots summarizing the effects of interventions on SAR modelled water table levels; higher values indicate wetter conditions. DF – ditch filling; DY – area enclosed by dyke; LG – lowering (scraping) of the ground; PO – pool construction. The effect of interventions is compared to the relevant control (ditch C for ditches; Fig. 4 and control area for other interventions; Fig. 5). Number of interventions shown between brackets. In 2024 mini bunds were constructed around interventions; the effects for these interventions are shown separately.

## 6. Discussion

The Sentinel-1 SAR data are well suited for the impact assessment of conservation interventions in wetlands such as the Boeung Prek Lapouv area in the Mekong delta. During the dry season the SAR data responded to variations in groundwater levels (measured through dipwells) and during the wet season to the extent of flooding. Separation of signals from dry and wet areas was essential for a meaningful analysis of the data. High noise levels in the SAR data hamper to some extent the identification of the start of a cycle (Adeli, 2020). The separate dry and wet-season SAR signals allowed for a cost-effective way to assess the impact of interventions on water retention across the region for longer time periods. This method is expected to save financial and human resources in data collection for intervention assessment over the future years.

The results indicate that the interventions carried out in 2022–2024 were by and large successful. The rank of the interventions from the most to the least impact was: 1) digging pools, 2) lowering ground, 3) blocking ditches and 4) building a dyke (Fig. 6; size of interventions in Section 2). Interestingly, in 2024, the soil excavated was used to surround the pools and the lowered grounds areas with mini bunds; the results suggest that these features result in a higher retention of water since they are wetter than those constructed in previous years (Fig. 6). On the downside, mini bunds make it harder to flush out the acidic water just after construction (Tucker and D’Abramo, 2008).

Construction of ephemeral pools, here the most impactful of all the interventions carried out, was found to have a positive impact on wetland restoration in other studies (Larson et al., 2016). Follow-on studies of the physical, chemical and biological parameters of these pools are planned to assess their long-term impact on fauna and flora (Kolozsvary & Holgeron, 2016). Lowering grounds and surrounding these by mini bunds was found to have the second largest impact. Scraping to remove the drier topsoil may have additional benefits e.g. in removing invasive species such as *Minosa pigra* and promoting the growth of a diverse wetland species community (Dalrymple et al 2003). Ditch filling is a relatively cheap intervention and was found to have the next largest effect promoting wet conditions. An important consideration is that the infill needs to withstand traffic from large animals such as cows and buffalo. Monitoring for at least a year is recommended to assess its true impact (Holden et al., 2016). The dyke embankment worked well in the 2024 dry season that followed the intervention but did not have a noticeable effect in the year after (Fig. 5). Other studies reported both positive impacts (Damm, 2013) and negative ones (Lien et al. 2022) as well. The dyke required further repairs when a 2-meter wide hole was formed by a fast-incoming flood in July 2024. Sluices that allow water to move more quickly would avoid this, but are difficult to maintain because of the remote location. Future studies will investigate the long-term effect of this dyke on groundwater retention.

The present study confirms that human-led restoration techniques using heavy machinery improve water retention in a protected landscape in the short term and may be a way to mitigate the effects of climate change long-term (Khanal et al. 2021). Sentinel-1 data were very successful in monitoring soil moisture and flooding and allowed comparison of conditions before and after interventions were made. Their use could be extended to other applications that require information on soil water saturation and that at present rely on regional proxies such as rainfall and extent of area flooded; this would improve assessments of flood risk and risk of landslides and thereby protect infrastructure and human lives.

As we face the limits of our freshwater use boundary, wetlands must be protected as the source and reservoir of freshwater needed for human and wildlife benefits. It is a priority to assure that the limited resource for protecting these key habitats is well spent. This study hopes to contribute to the welfare of wetlands and people alike.

## Acknowledgements

We are grateful to the WWT Cambodia team for their support during project implementation and to the Field Monitoring Team Mr Khann and Mr Panha. We sincerely thank the Ministry of Environment, the Ministry of Land Management, Urban Planning and Construction; and His Excellency Srey Sunleang, Director General of the General Directorate of Natural Protected Areas for their support. We extend our gratitude to Mr. Hong Chamnan, Director of the Department of Freshwater Wetlands Conservation, and to Mr. Visal Yoeung, Chief of Office for their invaluable assistance. Furthermore, we thank Mr Socheat and Mr Sarim, the BPL ranger, Mr Monly, Mr Phearak and Mr Vath, the Provincial Department of Environment of the region of Takeo, His Excellency Mr Ouch Phea, the Takeo governor and the deputy governor for their support. In addition, we warmly thank the 19 village chiefs, the 6 commune chiefs and the 2 district heads for their approval of this project.

This work was supported by the Critical Ecosystem Partnership Fund (CEPF-111877), the Fondation L’Occitane, the Fondation Prince Alberto II de Monaco, the Disney Conservation Fund, and WWT.

Conceptualization: All; Data curation: PS; Formal analysis: SOL, DK, BRS; Funding acquisition: BRS, DK, GMH; Methodology: SOL; Project administration: DK, BRS; Resources: GMH, OvB; Software: SOL; Supervision: GMH, OvB, DK; Validation: SOL; Visualization SOL; Writing – original draft: SOL, DK, BRS. Writing – review & editing: All.

The datasets (groundwater levels, locations of dipwells and interventions, 30-m DEM, processed SAR data, R code to estimate flooding) are available on the Open Science Frameworks: https://osf.io/cj5nd/?view_only=5d8aca511ffc45d2ad4f14da372e54df.

## Bibliography

1. Adeli, S., Salehi, B., Mahdianpari, M., Quackenbush, L. J., Brisco, B., Tamiminia, H., and Shaw, S. (2020). Wetland Monitoring Using SAR Data: A Meta-Analysis and Comprehensive Review. Remote Sens., 12, 2190. doi: 10.3390/rs12142190.

2. Bangira, T., Iannini, L., Menenti, M., Van Niekerk, A. and Vekerdy, Z. (2021) Flood Extent Mapping in the Caprivi Floodplain Using Sentinel-1 Time Series. IEEE J. Sel. Top. Appl. Earth Obs. Remote Sens., 14, 5667–5683, doi: 10.1109/JSTARS.2021.3083517.

3. Bennett, N.J., 2016. Using perceptions as evidence to improve conservation and environmental management. Conserv. Biol., 30, 582–592. doi: 10.1111/cobi.12681.

4. BirdLife International (2016). Antigone antigone. The IUCN Red List of Threatened Species 2016: e.T22692064A93335364. doi: 10.2305/IUCN.UK.2016-3.RLTS.T22692064A93335364.en. Accessed on 15 May 2025.

5. Brooks, T.M., Wright, S.J. and Sheil, D., 2009. Evaluating the success of conservation actions in safeguarding tropical forest biodiversity. Conserv. Biol., 23, pp.1448–1457. doi: 10.1111/j.1523-1739.2009.01334.x.

6. Campbell, I.C. (2012). Biodiversity of the Mekong Delta. In: Renaud, F., Kuenzer, C. (eds) The Mekong Delta System. Springer Environmental Science and Engineering. Springer, Dordrecht. doi: 10.1007/978-94-007-3962-8_11.

7. Copernicus Climate Change Service (C3S), (2024): ERA5-Land monthly total precipitation from 1-2021 to 9-2024. Copernicus Climate Change Service (C3S) Climate Data Store (CDS), doi: 10.24381/cds.68d2bb30 (Accessed on 30-Oct-2024).

8. Copernicus Sentinel data (2025) obtained from the Copernicus Data Space Ecosystem. https://dataspace.copernicus.eu/. Downloaded in 2024 and 2025.

9. Eyler, B., 2019. Last days of the mighty Mekong. Bloomsbury Publishing. London, UK. pp 384.

10. Dalrymple G.H., Doren R.F., O’Hare N.K., Norland M.R. & Armentano T.V. (2003) Plant colonization after complete and partial removal of disturbed soils for wetland restoration of former agricultural fields in Everglades National Park. Wetlands, 23, 1015–1029.

11. Damm, C. (2013) Ecological restoration and dike relocation on the river Elbe, Germany. Scientific Annals of the Danube Delta Institute 19, 79.

12. Fernández, D., Kerhoas, D., Dempsey, A., Billany, J., McCabe, G. and Argirova, E., 2022. The current status of the world’s primates: Mapping threats to understand priorities for primate conservation. Int. J. Primatol., 43, 15–39. doi: 10.1007/s10764-021-00242-2.

13. Fernández-Urrutia, M., Arbelo, M., and Gil, A. (2023). Identification of paddy croplands and its stages using remote sensors: A systematic review. Sensors, 23, 6932. doi: 10.3390/s23156932

14. Ferraro, P.J. and Pressey, R.L., 2015. Measuring the difference made by conservation initiatives: protected areas and their environmental and social impacts. Philos. Trans. R. Soc. London, Ser. B, 370, p.20140270

15. González I., Déjean S. (2023). CCA: Canonical Correlation Analysis. R package version 1.2.2, CRAN.R-project.org

16. Hoffmann, M., Hilton-Taylor, C., Angulo, A., Böhm, M., Brooks, T.M., Butchart, S.H., Carpenter, K.E., Chanson, J., Collen, B., Cox, N.A. and Darwall, W.R., et al. (2010) The impact of conservation on the status of the world’s vertebrates. Science, 330, 1503–1509. doi: 10.1126/science.1194442.

17. Holden, Joseph, Sophie M. Green, Andy J. Baird, Richard P. Grayson, Gemma P. Dooling, Pippa J. Chapman, Christopher D. Evans, Mike Peacock, and Graeme Swindles. “The impact of ditch blocking on the hydrological functioning of blanket peatlands.” Hydrological Processes 31, no. 3 (2017): 525–539.

18. Indo-Burma Ramsar Regional Initiative (2022) Indo-Burma Wetland Outlook 2022, IUCN

19. Khanal, S., Lutz, A. F., Kraaijenbrink, P. D. A., van den Hurk, B., Yao, T., & Immerzeel, W. W. (2021). Variable 21st century climate change response for rivers in High Mountain Asia at seasonal to decadal time scales. Water Resour. Res., 57, e2020WR029266. doi: 10.1029/2020WR029266.

20. Kolozsvary, M. B., and M. A. Holgerson. “Creating temporary pools as wetland mitigation: how well do they function?.” Wetlands 36 (2016): 335–345.

21. Kornelsen, K. C. and Coulibaly, P. (2013) Advances in soil moisture retrieval from synthetic aperture radar and hydrological applications, J. Hydrol., 476, 460–489. doi: 10.1016/j.jhydrol.2012.10.044.

22. Larson, Danelle M., John Riens, Sheldon Myerchin, Shawn Papon, Melinda G. Knutson, Sara C. Vacek, Sarah G. Winikoff, Mindy L. Phillips, and John H. Giudice. “Sediment excavation as a wetland restoration technique had early effects on the developing vegetation community.” Wetlands Ecology and Management 28, no. 1 (2020): 1–18.

23. Li, X., Liu, J.P., Saito, Y. and Nguyen, V.L., 2017. Recent evolution of the Mekong Delta and the impacts of dams. Earth Sci. Rev., 175, 1–17. doi: 10.1016/j.earscirev.2017.10.008.

24. Lien, B.T.B., Ngan, N.T.T., Kumar, P., Dang, T.T.T., Hong, T.T.K., Ty, T.V., Avtar, R. and Minh, H.V.T., 2022. Assessing the impacts of dike systems on water quality in natural reserves of the Vietnamese Mekong Delta. Urban Science, 6, 21., doi:10.3390/urbansci6010021.

25. Minderhoud, P.S., Erkens, G., Pham, V.H., Bui, V.T., Erban, L., Kooi, H. and Stouthamer, E., 2017. Impacts of 25 years of groundwater extraction on subsidence in the Mekong delta, Vietnam. Environ. Res. Lett., 12, p.064006. doi: 10.1088/1748-9326/aa7146.

26. Muñoz-Sabater, J., Dutra, E., Agustí-Panareda, A., Albergel, C., Arduini, G., Balsamo, G., Boussetta, S., Choulga, M., Harrigan, S., Hersbach, H., Martens, B., Miralles, D. G., Piles, M., Rodríguez-Fernández, N. J., Zsoter, E., Buontempo, C., and Thépaut, J.-N. (2021) ERA5-Land: a state-of-the-art global reanalysis dataset for land applications, Earth Syst. Sci. Data, 13, 4349–4383. doi: 10.5194/essd-13-4349-2021.

27. Oldekop, J.A., Holmes, G., Harris, W.E. and Evans, K.L., 2016. A global assessment of the social and conservation outcomes of protected areas. Conserv. Biol., 30, 133– 141. doi: 10.1111/cobi.12568.

28. Pham-Duc, B., Papa, F., Prigent, C., Aires, F., Biancamaria, S., & Frappart, F. (2019). Variations of Surface and Subsurface Water Storage in the Lower Mekong Basin (Vietnam and Cambodia) from Multisatellite Observations. Water, 11, 75. doi: 10.3390/w11010075.

29. Pressey, R.L., Visconti, P., McKinnon, M.C., Gurney, G.G., Barnes, M.D., Glew, L. and Maron, M., (2021) The mismeasure of conservation. Trends Ecol. Evol., 36, 808– 821. doi: 10.1016/j.tree.2021.06.008.

30. R Core Team (2024). R: A language and environment for statistical computing. R Foundation for Statistical Computing, Vienna, Austria. https://www.R-project.org/.

31. Sabel, D., Naeimi, V., Greifeneder, F., Wagner, W. (2015). Investigating Radar Time Series for Hydrological Characterisation in the Lower Mekong Basin. In: Kuenzer, C., Dech, S., Wagner, W. (eds) Remote Sensing Time Series. Remote Sensing and Digital Image Processing, vol 22. Springer, Cham. 10.1007/978-3-319-15967-6_17.

32. Seng Kim Hout (2004) Management of the Boeung Prek Lapouv Sarus crane conservation area, Takeo Province, MSc. Thesis, Prek Leap Agricultural University, Phnom Penh, Cambodia.

33. Sivuth, S, (2021) Wildfowl & Wetlands Trust (WWT) aerial survey of Anlung Pring and Boeung Prek Lapouv protected landscapes in Kampong Trach District, Kampot Province and Koh Andaet and Borei Cholsar District, Takeo Province, Cambodia, Aruna Technology Ltd, WWT Slimbridge, United Kingdom. 22 pp.

34. SNAP (2024) -ESA Sentinel Application Platform v10.0.0, http://step.esa.int x(accessed Sep. 2024).

35. Sophanna, L., H. Pok, and T. Avent (2019) Climate Change Vulnerability Assessment for Boueng Prek Lapouv Protected Landscape, Cambodia. Bangkok, Thailand: IUCN ARO. X + 32 pp.

36. Tucker C.S. and D’Abramo L.R. (2008) Managing high pH in freshwater ponds. Southern Regional Aquaculture Cente, Mississippi State University Stoneville, Mississippi (USA).

37. WWT (2023a) The Status of Wetland Habitats in the Cambodia Lower Mekong Delta. WWT, Slimbridge, UK.

38. Yang, H., Liu, D., Xiao, H. et al. (2025) Better plans are needed for mitigating the ecological impacts of Cambodia’s Funan Techo Canal. Nat. Ecol. Evol. 9, 185–186. doi: 10.1038/s41559-024-02605-3

