## Supplement for "Assessing wetland conservation action effectiveness in the Mekong Delta with Sentinel-1 Synthetic Aperture Radar data"

### S.1 Selection criteria to identify areas suitable for restoration

- Elevation. Data (from the July 2021 UAV aerial survey) used to identify low lying areas across BPL. Areas below 2.40 m a.s.l. preferred.
- Land-use. 2020 land-use map used.
    - 'grassland', 'tall grass' and 'scrub' selected.
    - 'wet grassland', 'aquatic vegetation', 'open water' excluded.
- Satellite imagery. Google Earth historical images accessed to identify areas of grassland that over the last 10 years showed visible signs of drying out e.g. invaded by scrub species
- Avoidance of channels used by local people to access their rice farms or any area known to be important to their livelihoods.
- Avoidance of land under ownership dispute.
- Avoidance of land near villages or individual rice farms. Advise provided by government.
- Avoidance of altering any natural water course.
- Avoidance of areas used by species of conservation concern eg Yellow-breasted bunting; Sarus crane roost or foraging areas.
- Avoidance of areas earmarked for flooded forest restoration.
- For ephemeral pools, avoidance of areas where soil data suggests the iron sulphide layer is less than 10 cm from the surface.

**Table S.1 Description of restoration activities 2022-2024**

| Intervention type | Reason | Construction technique | Output (2022-2024) |
|---|---|---|---|
| Block human-made ditches / Infill human-made ditches | To reduce the loss of water (through drainage) at the end of the flood season; raise groundwater levels within approximately 5m either side of the ditches; provision of open water (blocked ditches only, not infilled) as foraging habitat for waterbirds including Sarus crane | Earth placed at strategic locations inside ditch channels to form an effective dam; typically 5m long. The earth is compacted by an excavator. In situations where excess spoil was generated from landscaping works, sections of ditch were fully infilled. | 60 sections of ditch blocked; total length = 4,080m<br><br>10 sections of ditch infilled; total length = 2,893m |
| Scraped ground / bunded scraped ground | Raise groundwater levels to encourage the establishment of 'wet grassland', in particular *Eleocharis dulcis*, the tubers of which are a known food source of Sarus crane. | • Vegetation layer (including roots) scraped off using a dozer.<br>• Approx. 10cm of the upper soil layer removed using a dozer and / or excavator.<br>• Central portion lowered up to 15cm to encourage flood water to drain towards the centre.<br>• Where possible, spoil used to construct a perimeter bund around the lowered ground. Bund approx. 2m (H) x 3m (wide at base). The earth is compacted by an excavator. | Scraped ground at 8 locations; total area = 63.84ha; 32.56 ha with perimeter bund |
| Ephemeral pools / Bunded ephemeral pools | Create habitat to encourage the establishment of water lily Nymphaea sp. The tubers are a known food source for Sarus crane in the region.<br><br>Benefit aquatic invertebrate and plant communities. | • Vegetation layer (including roots) scraped off using a dozer.<br>• Approx. 30cm of the upper soil layer removed using a dozer and / or excavator. Shallow sloping margins. Deeper central areas (up to 40cm) to retain flood water for longer.<br>• Where possible, spoil used to construct a | 10 ephemeral pools; total surface area = 18.96ha; 17.55ha with perimeter bund |

| | | | |
|---|---|---|---|
| | | perimeter bund around the pond boundary. Bund approx. 2m (H) x 3m (wide at base).<br>• Nymphaea plants transplanted into the pools. | |
| Dyke impoundment | To retain surface water and raise groundwater levels for longer periods of time at the end of the flood season. Expected to create a range of habitats attractive to waterbirds including 'wet grassland' and water lily. | A 2m (H) x 5m (wide at base) earth bund constructed using multiple excavators. Earth was obtained by excavating a ditch (approx. 2m wide x 1m deep) inside the enclosure, from other restoration landscaping activities, and suitable pockets of soil close to the dyke. The earth is compacted by an excavator. | 3,713m dyke wall; total enclosed area = 83.80ha |

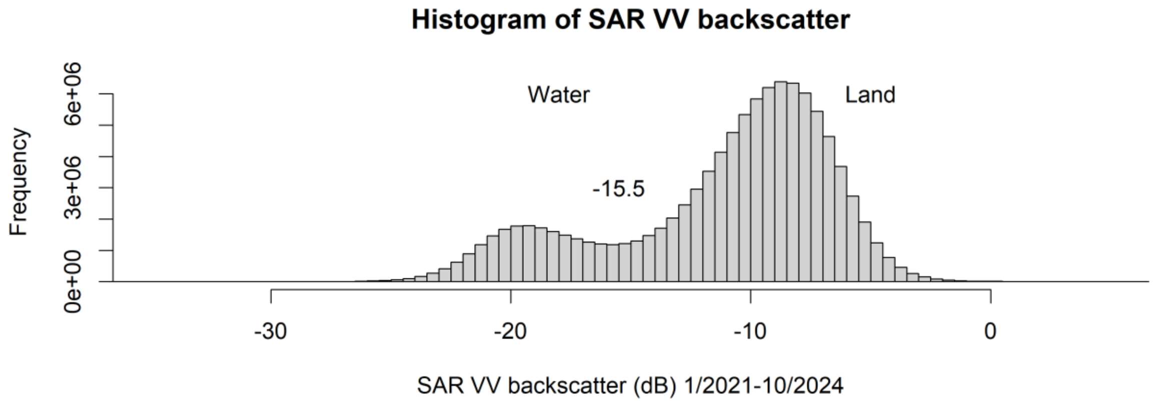


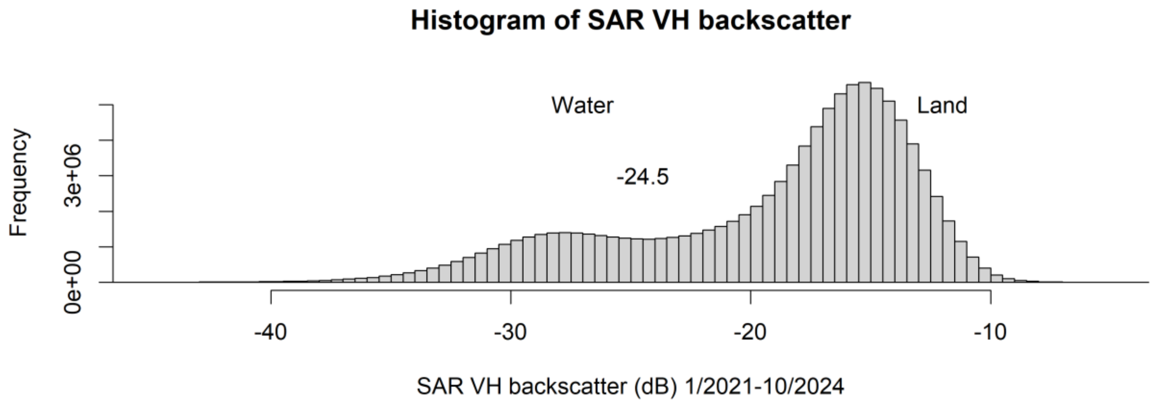


*Figure S.1: Histograms of SAR backscatter (dB). Top: SAR VV backscatter. Bottom: SAR VH backscatter. Numbers (-15.5 and -24.5) indicate the thresholds used to identify flooding events.*

### S.2 Details of the algorithm to separate time series from dry and flooded conditions

The algorithm proposed in the present study is similar in spirit to the one developed by Bangira et al. (2021). It groups consecutive observations and attributes a cycle of decreasing and increasing values to either a flooding cycle or a drying cycle. Time series of VV, VH and the ratio between them, VH / VV, are converted to dB (by taking the log and multiplying by 10; note that VH / VV becomes 10 * log VH – 10 * log VV). The subsequent analysis consists of the following steps:

1. The signal in VV, VH and VH/VV (in dB) is smoothed by averaging over a window of 5 time steps (- 1 month to + 1 month).
2. A cycle starts with a decline, which is identified by a first temporal derivative (change from the previous time step) lower than -0.1 dB in at least two of the smoothed VV, VH or VH/VV time series.
3. A decline is removed if its neighbours on either side are not a decline; likewise, if a point is not declining but its two neighbours are it is identified as a decline.

4. Consecutive sets of declining points are assigned to a group, for each group it is established if the decline is caused by flooding or by a decrease in soil moisture. The decline is attributed to a flood if one of the following criteria is met:
    a. Any VV is below -15.5 dB, any VH below -25.5 dB or any VH/ VV is below -9 dB.
    b. The start of the decline coincides with a wet spell (monthly precipitation above 50 mm). For months with precipitation reanalysis missing (Oct – Dec 2024) the monthly averages of 2021 – 2023 were used in the analysis.
5. Once a declining group is attributed to either a dry event or a flooding event the upward trajectory is added. The upward trajectory continues as long as 2 out of three (VV, VH or VH/VV) time series have values below the respective median of the downwards trajectory. From this point, the upward trajectory is further extended as long as two out of three (VV, VH or VH/VV) time series show an increase.

R code used to identify flooded conditions:

```
function(Year, vv, vh, ppn, per=24){

  #Year  - year as decimal, e.g first twice-monthly composite of 2022 is
  #2022 + 0.5/24, second composite 2022 + 1.5/24 etc.
  #vv    - twice monthly time series of SAR vv data
  #vh    - same but for vh data
  #ppn   - twice monthly precipitation time series (may need to be
  #        adjusted to vv and vh timestep)
  #per   - number of vv and vh timesteps in a year;
  #        assumed twice monthly data – 24 time steps per year

  # set thresholds to identify flooded conditions – adjust as needed
  vh.thr <- -24.5 # estimated from histograms
  vv.thr <- -15.5 # same
  vhvv.thr <- -9  # estimated from analysis transect A-C time series
  flooded <- rep(NA, length(vv)) # create empty vector
  ny <- ceiling(length(vv)/per)

#  convert to decibel if needed
  if (mean(vv, na.rm=T) > 0) vv <- 10*log10(vv)
  if (mean(vh, na.rm=T) > 0) vh <- 10*log10(vh)

# calculate moving averages
  tmp <- c(NA, NA, vv, NA, NA)
  vv.ts <- apply( cbind(tmp[-c(1:4)],
                        tmp[-c(1:3,length(tmp))],
                        tmp[-c(1:2,(0:-1) + length(tmp))],
                        tmp[-c(1,(0:-2) + length(tmp))],
                        tmp[-c( (0:-3) + length(tmp))]),1,mean,na.rm=T)

  d.vv.ts <- c(0, diff(vv.ts)) # first derivative indicates increase or
decrease of vv.ts

  tmp <- c(NA, NA, vh, NA, NA)
  vh.ts <- apply( cbind(tmp[-c(1:4)],
                        tmp[-c(1:3,length(tmp))],
                        tmp[-c(1:2,(0:-1) + length(tmp))],
                        tmp[-c(1,(0:-2) + length(tmp))],
                        tmp[-c( (0:-3) + length(tmp))]),1,mean,na.rm=T)

  d.vh.ts <- c(0, diff(vh.ts))
  vh.vv.ts <- vh.ts - vv.ts # ratio vh and vv in dB
  d.vh.vv.ts <- c(0,diff(vh.vv.ts))
```

```
  # check if sufficient non-missing data
  if (sum(is.finite(vh.vv.ts)) > (0.9*length(vh.vv.ts)) ){

    #search for start of cycles / adjust threshold -0.1 as needed
    declining <- (apply(cbind(d.vh.vv.ts, d.vh.ts, d.vv.ts) < -0.1,1,sum,
na.rm=T) >= 2) & !is.na( d.vh.vv.ts)

    # remove false declines or increases; identify by checking if
    # neighbours do something different (noisy data)

    for( i in 2:(length(declining)-1) ){
      if ( (declining[i] ) & !declining[i+1] & !declining[i-1] )
declining[i] <- declining[i-1]
    }

    ld <- length(declining)

    # further checks on neighbours
    sum.decl <- sum(declining, na.rm=T)
    p.sum <- 0
    while (p.sum < sum.decl){
      for( i in 2:(length(declining)-1) ){
          tmp.sum <- sum((c(d.vh.vv.ts[i-1], d.vh.ts[i-1], d.vv.ts[i-1]) <
0), na.rm=T )
          if ( declining[i] & !declining[i-1] & ( tmp.sum >=2) )
declining[i-1] <- TRUE
          tmp.sum <- sum((c(d.vh.vv.ts[i+1], d.vh.ts[i+1], d.vv.ts[i+1]) <
0), na.rm=T )
          if ( declining[i] & !declining[i+1] & (tmp.sum >=2) )
declining[i+1] <- TRUE
      }
      p.sum <- sum.decl
      sum.decl <- sum(declining, na.rm=T)
    }

    for( i in 2:(length(declining)-1) ){
      if ( (!declining[i] ) & declining[i+1] & declining[i-1] )
declining[i] <- declining[i-1]
    }

    # check ends
    if(declining[1] != declining[2]) declining[1] <- declining[2]
    if(declining[ld] != declining[ld-1]) declining[ld] <- declining[ld-1]

    #start looking for increasing part after decline, idenitify groups of
declining timesteps
    groups <- rep(0, length(declining))
    ig <- 1
    if(declining[1]) groups[1] <- ig
    for( i in 2:length(declining) ){
      if (declining[i] & declining[i-1]){
         groups[-1:0+i] <- ig
      }
      else if (declining[i] & !declining[i-1]) {  # new group, increase
counter
        ig <- ig + 1
      }
    }
    ind <- 1:length(groups)
    u.groups <- sort(unique(groups))[-1]
```

```
    for (i in u.groups){
      t.ind <- ind[groups==i]
      min.vh.vv <- min( vh.ts[t.ind] , na.rm=T)
      i.m <- ceiling(length(t.ind)/2)
      x.ind <- sort( unique( c(t.ind[1:i.m] - 4, t.ind) ) )
      x.ind <- x.ind[x.ind > 0]

      # groups need to be sufficiently long, flooding is identified if
start decline has large ppn
      # or if some values in the time series are indicative of flooded
conditions

      if( ( (min(ppn[x.ind], na.rm=T) < 50) | (length(t.ind) < 3) ) &
           !( (length(t.ind) > 8) | (any(vh.ts[t.ind] < vh.thr, na.rm=T) )
) ){
        declining[t.ind] <- FALSE
        groups[t.ind] <- 0
      }
    }
    u.groups <- sort(unique(groups))[-1]

    flooded <- declining
    dry.thr <- quantile( vh.ts[declining], p=0.95, na.rm=T)

 # find increasing part of cycle
    for (i in u.groups){
      t.ind <- ind[groups==i]
      m.vh <- median(vh.ts[t.ind], na.rm=T)
      m.vv <- median(vv.ts[t.ind], na.rm=T)
      m.vhvv <- median((vh.vv.ts)[t.ind], na.rm=T)

      t.iii <- iii <- min(c(t.ind[length(t.ind)]+1, length(groups)),
na.rm=T)
# fill up increasing values until the median of the declining values
      while (iii <= length(groups)){
        tsum <- sum ( c( vh.ts[iii] < m.vh, vv.ts[iii] < m.vv,
vh.vv.ts[iii] < m.vhvv), na.rm=T)
        if (tsum >=2){
          flooded[iii] <- TRUE
          t.iii <- iii <- iii + 1
        }
        else{
          iii <- length(groups)+1
        }
      }

# now fill up while increase continues
      iii <- t.iii
      while (iii <= length(groups)){
        tsum <- sum ( c( vh.ts[iii] > vh.ts[iii-1], vv.ts[iii] > vv.ts[iii-
1],
                         vh.vv.ts[iii] > vh.vv.ts[iii-1]), na.rm=T)
        if (tsum >=2){
          flooded[iii] <- TRUE
          iii <- iii + 1
        }
        else{
          iii <- length(groups)+1
        }
      }
```

```
      flooded[(vh.ts > dry.thr) & !is.na(vh.ts)] <- 0
    }   # end for
  } # end if
  list(ma.vv=vv.ts, ma.vh=vh.ts, flood=flooded )
} # end of function
```

Bibliography / source of inspiration

Bangira, T., Iannini, L., Menenti, M., Van Niekerk, A. and Vekerdy, Z. (2021) Flood Extent Mapping in the Caprivi Floodplain Using Sentinel-1 Time Series. IEEE J. Sel. Top. Appl. Earth Obs. Remote Sens.14, 5667-5683, doi: 10.1109/JSTARS.2021.3083517.
